# Divergent Strategies of Mycorrhiza-Mediated Drought Adaptation in Poplar

**DOI:** 10.1101/2025.10.01.679871

**Authors:** Huili Shi, Zhuchou Lu, Andrea Polle

**Author notes:** permanent address: Research Institute of Subtropical of Forestry, Chinese Academy of Forestry, Hangzhou, Zhejiang, 311400, P. R. China. The author responsible for distribution of materials integral to the findings presented in this article in accordance with the policy described in the Instructions for Authors (https://academic.oup.com/plphys/pages/General-Instructions) is Andrea Polle.

## Abstract

Mycorrhizal symbiosis shapes plant growth and stress resilience. Here, we compared physiological and molecular responses of poplars colonized by *Paxillus involutus* (Pi) or *Cenococcum geophilum* (Cg) under control conditions, drought stress, and recovery. Both fungal species primed distinct local (root) and systemic (leaf) defenses compared to non-inoculated (Ni) plants. Cg-colonized poplars exhibited constitutively elevated transcripts of heat shock proteins (*HSP*s), galactinol synthase, and aquaporins in roots and leaves, irrespective of drought. Pi colonization enhanced growth and nitrogen-use-efficiency along with transcriptional increases of TOR/RAPTOR complex. Under severe soil moisture decline, Pi and Ni poplars showed reduced water potential, photosynthesis, growth, and leaf shedding, whereas Cg-colonized plants maintained water status, sustained photosynthesis, and retained foliage. These results reveal two contrasting mycorrhiza-mediated drought strategies in poplar: Pi fosters stress acclimation via drought-induced leaf abscission, enabling rapid recovery; Cg suppresses growth even without stress, conferring constitutive tolerance. Ectomycorrhizal species thus occupy different positions on the growth– defense trade-off spectrum. Such species-specific effects have important ecological and applied implications, enabling targeted use of EM fungi in forestry and agriculture depending on whether maximizing productivity or enhancing stress resilience is the primary goal.

## Introduction

Over the past two decades, episodes of severe drought have led to extensive tree mortality in forests worldwide (Allen et al., 2010; Choat et al., 2018). Climate change is projected to further intensify drought severity on a global scale (McDowell et al., 2020). Prolonged dry conditions are driving pronounced declines in local water tables (Williams et al., 2022) with detrimental impacts on both mature and juvenile trees (Moser et al., 2010; Subedi et al., 2021). Riparian species such as poplars (*Populus* spp.) are particularly vulnerable to these conditions (Garssen et al., 2014) due to their inherently water-spending life strategy (Polle et al., 2019).

The decline of wild and cultivated poplar populations is of considerable concern, as these trees are keystone components of riparian ecosystems (Cronk, 2005; De Carvalho et al., 2010; Kõrkjas et al., 2021) and hold high economic value as biomass crops in short-rotation forestry (Taylor et al., 2019).

Among the strategies for improved tree protection under global change, the use of beneficial microbial inoculations, especially mycorrhizal fungi, has emerged as a promising strategy for enhancing tree resilience (Groover et al., 2025). Under natural field conditions, poplar roots typically form symbioses with ectomycorrhizal (EM) fungi, which can enhance host tolerance to abiotic stress (Dreischhoff et al., 2020; Rosenkranz et al., 2023). Nevertheless, current knowledge of the molecular mechanisms and physiological responses governing poplar drought tolerance in association with EM fungi remains limited. Addressing this knowledge gap is of high priority, since growth physiology and nutrient acquisition are likely key determinants of drought resistance in trees (Gessler et al., 2017).

Emerging evidence suggests that different ectomycorrhizal (EM) fungi confer distinct physiological benefits to their host plants. For example, *Populus* spp. colonized by either *Paxillus involutus* or *Laccaria bicolor* exhibit pronounced differences in leaf gas exchange, carbon allocation, nutrient supply, and biomass production (Shinde et al., 2018; Shi et al., 2024). Bouffaud et al. (2020) reported differential transcriptional responses in oak (*Quercus*) seedlings associated with *P. microcarpus*, *P. involutus*, or *L. bicolor*, although the physiological consequences of these differences remain unclear (Bouffaud et al., 2020).

EM symbioses may facilitate water uptake by increasing root hydraulic conductivity or water transport capacity through aquaporin-mediated processes (Lehto and Zwiazek, 2011). However, this ability appears to depend on the particular combination of plant and fungal species. For instance, *Hebeloma cylindrosporum* improved root hydraulic conductivity compared with other fungal symbionts (Bogeat-Triboulot et al., 2004; Siemens and Zwiazek, 2008). *L. bicolor* induced higher hydraulic conductivity and greater aquaporin expression in white spruce (*Picea glauca*)) (Xu et al., 2015), but not in trembling aspen (*Populus tremuloides*) (Xu et al., 2016). Conversely, some studies have reported either no change or even reductions in aquaporin gene expression under drought in EM plants colonized by *P. involutus* (Danielsen and Polle, 2014) or *L. bicolor* (Calvo-Polanco et al., 2019).

Among the EM fungi, *Cenococcum geophilum* stands out for its drought tolerance (Peter et al., 2016). This abundant generalist forms ectomycorrhizas with numerous tree species, including poplars (Bahram et al., 2011). Field surveys and controlled experiments have demonstrated that *C. geophilum* survives more effectively in dry environments than many other EM fungi (Pigott, 1982; di Pietro et al., 2007; Kipfer et al., 2010), a few exceptions have been reported (Nilsen et al., 1998). Mechanistically, its drought tolerance has been linked to increased expression of fungal genes associated with fatty acid metabolism, peroxisomal reactive oxygen species (ROS) production, and terpenoid biosynthesis (Li et al., 2022). A hallmark of *C. geophilum* symbiosis is the strong upregulation of fungal aquaporins compared with free-living mycelium (Peter et al., 2016), suggesting an enhanced capacity for water transfer to the host.

Other EM fungi employ different mechanisms to influence plant drought performance. For example, *P. involutus* forms rhizomorphs that can facilitate water transport (Agerer, 2001) and may enhance drought protection in poplar (Beniwal et al., 2010). *L. bicolor* triggers transcriptomic changes in roots, but under moderate drought stress, no physiological differences were detected between colonized and non-inoculated poplars (de Freitas Pereira et al., 2023). However, it remains unknown whether, and by what mechanisms, root colonization by *C. geophilum* or *P. involutus* preconditions their host to withstand drought. A key question is whether different EM fungal species elicit similar or distinct beneficial molecular and physiological processes in their hosts to improve drought resilience

In this study, we compared the molecular and physiological performance of grey poplar (*Populus × canescens*) colonized by either *P. involutus* (Pi) or *C. geophilum* (Cg) with that of non-inoculated plants (Ni). Trees were grown under controlled greenhouse conditions and subjected to a long-term, severe drought treatment, followed by re-watering to examine recovery. We hypothesized that (1) mycorrhizal poplars would exhibit stress-priming effects, irrespective of the associated EM species, thereby increasing drought resistance compared with Ni plants. Alternatively, we expected that (2) symbioses with different EM fungi would enhance either plant growth or stress tolerance, resulting in a trade-off between biomass production and defense activation. Since plants with greater biomass are likely to consume more water, we further considered that (3) differences in plant damage could arise from EM-induced growth stimulation. To address these hypotheses, we measured soil water content, plant growth, nitrogen concentration, predawn water potential, and transcriptomic changes in roots and leaves under well-watered, drought-stressed, and re-watered conditions. Transcriptomic results were validated with Cg-colonized and non-mycorrhizal poplar plantlets under sterile conditions.

## Results

### Photosynthesis and pre-dawn water potential of Cg poplars are unaffected by drought in contrast to Pi and Ni poplars

Drought stress was imposed by 80% reduction of water supply. Compared with the well-watered plants (mean soil moisture 0.237 ± 0.005 m³ m^-^³, Fig. 1), reduced water supply caused a gradual decrease in soil moisture, until mean levels of 0.073 ± 0.009 m³ m^-^³ were reached (Fig. 1). Thereafter, we kept the soil moisture constant at this level (Fig. 1). After rewatering, the soil moisture recovered within one day to levels close to those of the well-watered plants (Fig. 1).

**Figure 1.**
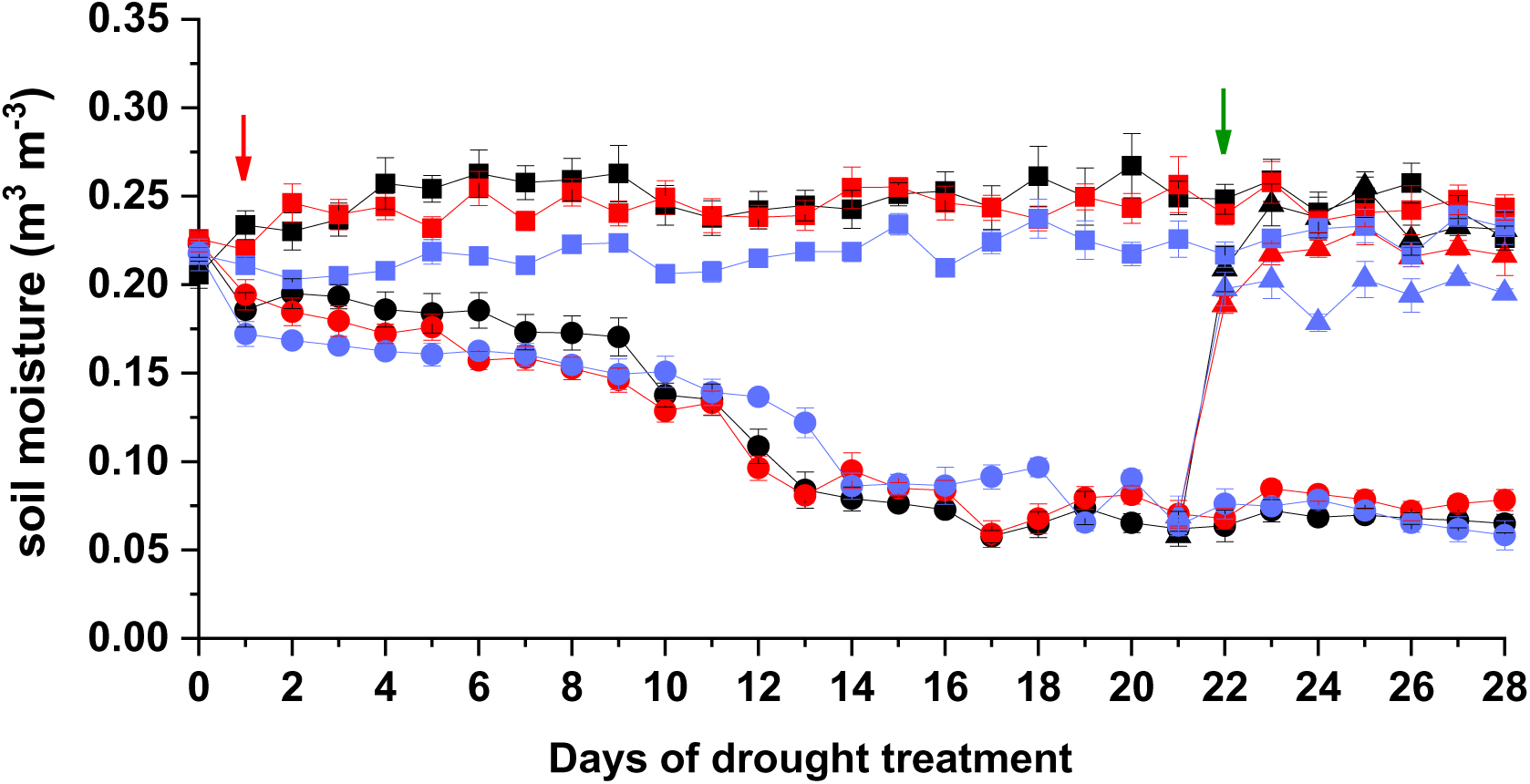
Dynamics of soil moisture in pots with mycorrhizal poplars during four experimental weeks before harvest. Plants were grown for 16 weeks under well-watered conditions and then divided in treatment groups: drought (circle), rewatered after three weeks of drought (triangles) and well-watered (square). The plants were colonized by *Paxillus involutus (*Pi: red), Cg: *Cenococcum geophilum (*Cg: blue) or were not inoculated (Ni: black). Data indicate means ± SE (n = 5 to 6 pots per treatment). The red arrow marks the start of the drought treatment, and the green arrow marks the start of the rewatering treatment.

Well-watered Ni, Pi, and Cg poplars showed no significant differences in photosynthesis (Fig. 2A), stomatal conductance (Fig. 2B) or pre-dawn water potential (Fig. 2C). Drought-stressed Ni and Pi plants had about 5-fold lower stomatal conductance and 3-fold lower photosynthesis rates than drought-exposed Cg poplars (Fig. 2A,B) and exhibited a drastic decline in the pre-dawn water potential, which was greater for the Pi than the Ni plants (Fig. 2C). Notably, upon reduced water supply, the Cg plants maintained gas exchange and pre-dawn water potentials at levels similar to those of well-watered plants, although the soil moisture declined in a manner similar to that of the Pi and Ni plants (Fig. 1). After rewatering, photosynthesis and stomatal conductance of Ni and Pi poplars recovered but did not reach the pre-drought values (Fig. 2A, B), despite complete recovery of the pre-dawn potential (Fig. 2C).

**Figure 2.**
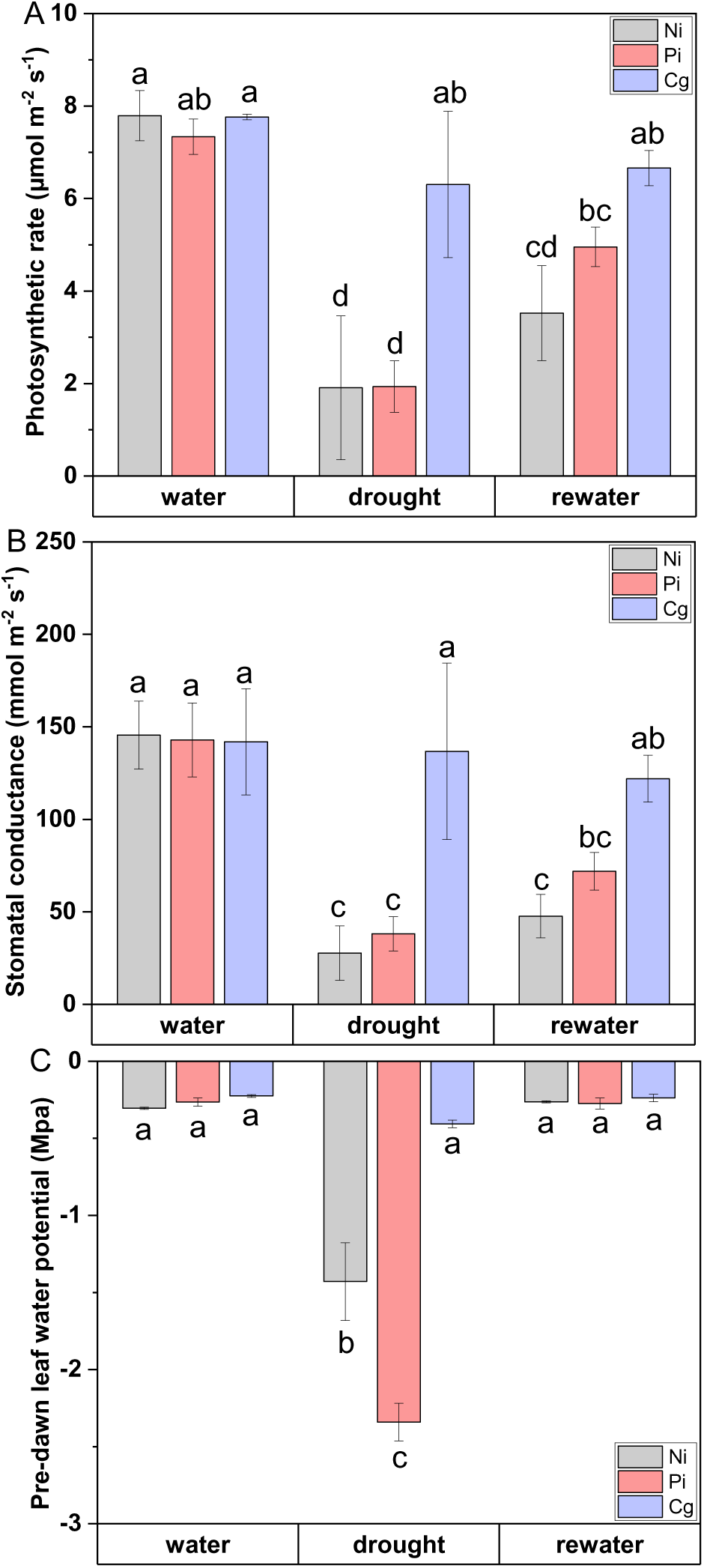
Photosynthesis (A), stomatal conductance (B), and pre-dawn leaf water potential (C) of mycorrhizal poplars. Data were measured during the last 2 weeks before harvest of well-irrigated and drought-stressed plants, and in the last week of rewatering treatment. Data indicate means ± SE (n= 3 to 4). Significant differences at *p* ≤ 0.05 between different treatments are indicated by different letters (Two-way ANOVA and post-hoc Fisher test). Ni: non-inoculated plants, Pi: *P. involutus*, Cg: *C. geophilum*, water: well-watered, drought, rewater: re-watered.

### Cg and Pi poplars exhibit high mycorrhization under stressed and non-stressed conditions

We tested whether the differences in plant physiology were related to differences in EM colonization rates. The EM colonization of Pi and Cg poplar root tips were in a similar range (55% to 68%, Table 1). Ni plants were not completely non-mycorrhizal because EM infections cannot be entirely avoided during a 5-month growth period under non-sterile greenhouse conditions but the Ni plants showed significantly lower EM colonization rates (8% under well-watered and 17% under drought) than EM-inoculated plants (Table 1). The EM colonization rates of Pi and Cg plants increased slightly under drought and were drastically higher than those of Ni plants. This result precludes that a greater stress response of Pi than that of Cg or Ni plants (Fig. 2) was a direct consequence of lower mycorrhization rates.

**Table 1.**
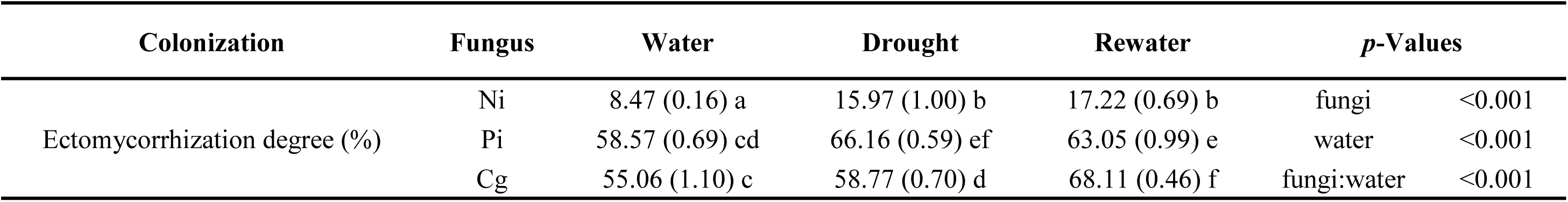
Mycorrhizal colonization (%) of vital poplar root tips. The plants were either non-inoculated (Ni) or inoculated with *P. involutus* (Pi) or *C. geophilum* (Cg) and cultivated under well-watered (water), drought-stressed (drought), or rewatered (rewater) conditions. Data indicate means (SE) of n= 5 to 6 plants per treatment. Data were processed with a beta regression model and ‘tukey adjusted comparison’ was applied as post test. Significant differences at p ≤ 0.05 are indicated by different letters. Calculated P-values listed in the last column indicate the comparison between fungal treatments (fungi), water treatments (water), and the interaction effect of fungal and water treatments (fungi:water). Ni: non-inoculation, Pi: *Paxillus involutus*, Cg: *Cenococcum geophilum* Fr., water: well-watered, rewater: re-watered.

### Pi stimulates biomass production under non-stressed conditions but leads to massive leaf shedding under drought, in contrast to Cg

Well-watered Pi plants had the greatest height growth rate (Fig. 3A). Stem diameter growth was also greatest for the Pi poplars, intermediate for Ni and lowest for the Cg poplars (Supplement Fig. S1). These growth differences corresponded to differences in whole plant biomass (Fig. 3B, Supplement Table S1) and in whole-plant N contents: Pi > Ni > Cg (Table 2). However, the weighted mean N concentration was greatest in Cg and lowest in Pi plants (Table 2). Similarly, the amount of N taken up per g gram of produced biomass (Supplement Fig. S2) showed Cg > Ni > Pi = 1.3 > 1.1 > 0.9 (p < 0.001 for each comparison). This indicates that N taken up by the Cg plants was less used for growth than that taken up by the Pi plants, implying that lower growth rates of the Cg plants were not caused by N limitation.

**Figure 3.**
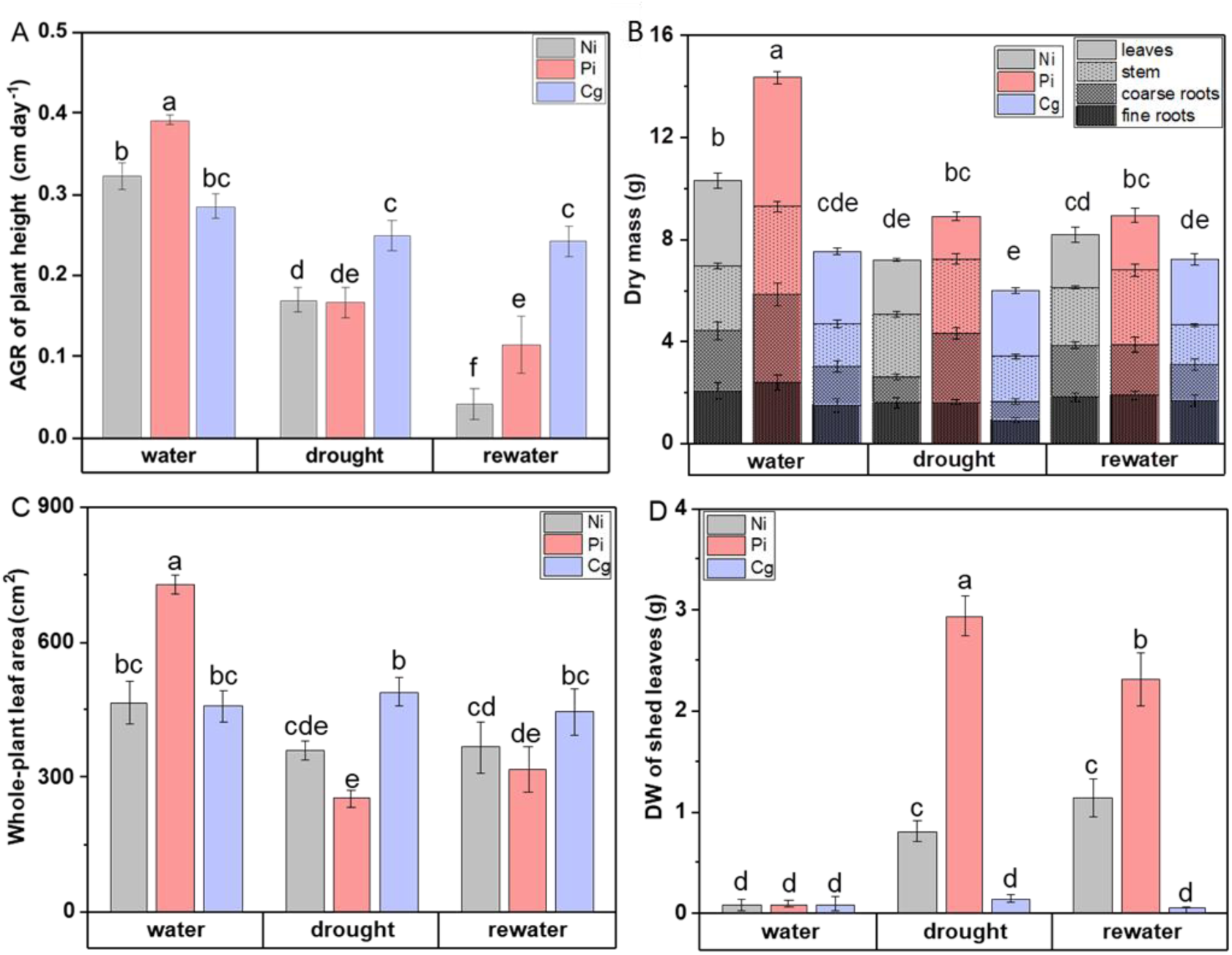
Growth rate (AGR) of plant height (A), whole-plant biomass (B), whole-plant leaf area (C) and dry mass of shed leaves (D) of mycorrhizal poplars in response to well-watered, drought and re-watering treatments. Data indicate means ± SE (n = 5 to 6 plant per treatment). In each panel, significant differences at *p* ≤ 0.05 are indicated by different letters (Two-way ANOVA and post-hoc Fisher test). DW: dry weight. Ni: non-inoculated, Pi: *Paxillus involutus*, Cg: *Cenococcum geophilum* Fr., water: well-watered (20 weeks), drought (4 weeks), rewater: re-watered (1 week).

**Table 2.** N content and weighted mean N concentrations of whole poplar plants. The plants were either non-inoculation (Ni) or colonized with *P. involutus* (Pi) or *C. geophilum* i(Cg). The plants well-watered (water), drought-stressed (drought) or rewatered (rewater). Data indicate means (SE) of n = 5 to 6 plants per treatment. Different letters indicate significant differences at p ≤ 0.05 (two-way ANOVA and post-hoc Fisher’s LSD test). Calculated P-values listed in the last column are indicate the comparison between fungal treatments (fungi), between water treatments (water), and the interaction of fungal and water treatments (fungi:water).

| Variable | Fungi | Water | Drought | Rewater | *p*-Values |  |
| --- | --- | --- | --- | --- | --- | --- |
| N content of the whole plant (mg) | Ni | 80.46 (8.21) b | 58.51 (1.92) cd | 65.00 (3.15) cd | Fungi | 0.009 |
|  | Pi | 102.09 (4.68) a | 58.66 (3.87) cd | 66.59 (3.72) cd | Water | <0.001 |
|  | Cg | 66.70 (3.44) cd | 54.55 (7.11) d | 71.16 (3.56) cd | fungi:water | <0.001 |
| Weighted mean N concentration (mg g<sup>-1</sup> DW) | Ni | 7.80 (0.33) cd | 8.16 (0.15) bc | 7.97 (0.28) cd | Fungi | <0.001 |
|  | Pi | 7.15 (0.15) de | 6.59 (0.20) e | 7.48 (0.43) cd | Water | 0.143 |
|  | Cg | 8.92 (0.33) ab | 9.13 (0.49) a | 9.81 (0.31) a | fungi:water | 0.300 |

Across the whole 4-week drought treatment, height and stem diameter growth were unaffected by drought in Cg poplars, whereas mean height growth rates of Pi and Ni poplars were about 2-fold lower than under well-watered conditions (Fig. 3 A). It should be noted that the growth rates of Ni and Pi plants declined dynamically during the drought period and were almost zero at the end of the drought period, whereas the height growth of Cg plants was unaffected (Fig. 3A). The height growth of Pi and Ni poplars recovered after rewatering (Fig. 3A). Pi poplars showed faster recovery of stem diameter growth than Ni poplars after rewatering (Supplement Fig. S1).

Whole-plant leaf area is decisive for plant water consumption and growth of poplar (Yu et al., 2019). In line with high growth rates, well-watered Pi poplars also exhibited a larger whole-plant leaf area than Ni or Cg plants (Fig. 3C). Significant leaf shedding was only observed during the drought phase (Fig. 3D). Pi plants exhibited more drastic leaf loss than Ni plants (Fig. 3D), resulting in severe biomass reduction (Fig. 3C). Only marginal leaf loss was observed for the Cg plants under drought (Fig. 3C). Although whole-plant leaf areas of Cg and Ni plants were similar under well-watered conditions, the Cg plants kept their leaf area under drought while that of the Ni plants declined because of leaf shedding (Fig. 3C, D).

In addition to biomass loss caused by leaf shedding, Pi plants showed reduction of the stem, the coarse, and the fine roots, while Ni plants showed a significant reduction of coarse root biomass under drought (Fig. 3B, Supplement Table S1). Cg plants showed no significant changes in biomass in response to drought (Fig. 3B, Supplement Table S1). However, under well-watered conditions, whole plant biomass of the Pi plants was about twice that of the Cg plants and after drought, the difference was shrunken to 1.5x (Fig. 3B, Supplement Table S1).

### Global transcriptomes differ in Pi and Cg-associated poplars under well-watered and stressed conditions

Roots and leaves of drought-stressed Pi plants showed the largest and Cg plants an intermediate number of DEGs (differentially expressed genes) compared with well-watered conditions (Fig. 4A, full list is shown in Supplement Table S2). Surprisingly, only three drought-induced DEGs were found in Ni leaves (Fig. 4A). Thus, mycorrhizal colonization strongly amplified drought-responsive transcriptional activity in a fungal species-specific manner. Re-watering resulted in a strong decrease in the number of DEGs in Pi and Cg plants, suggesting return to non-stressed conditions (Fig. 4A).

**Figure 4.**
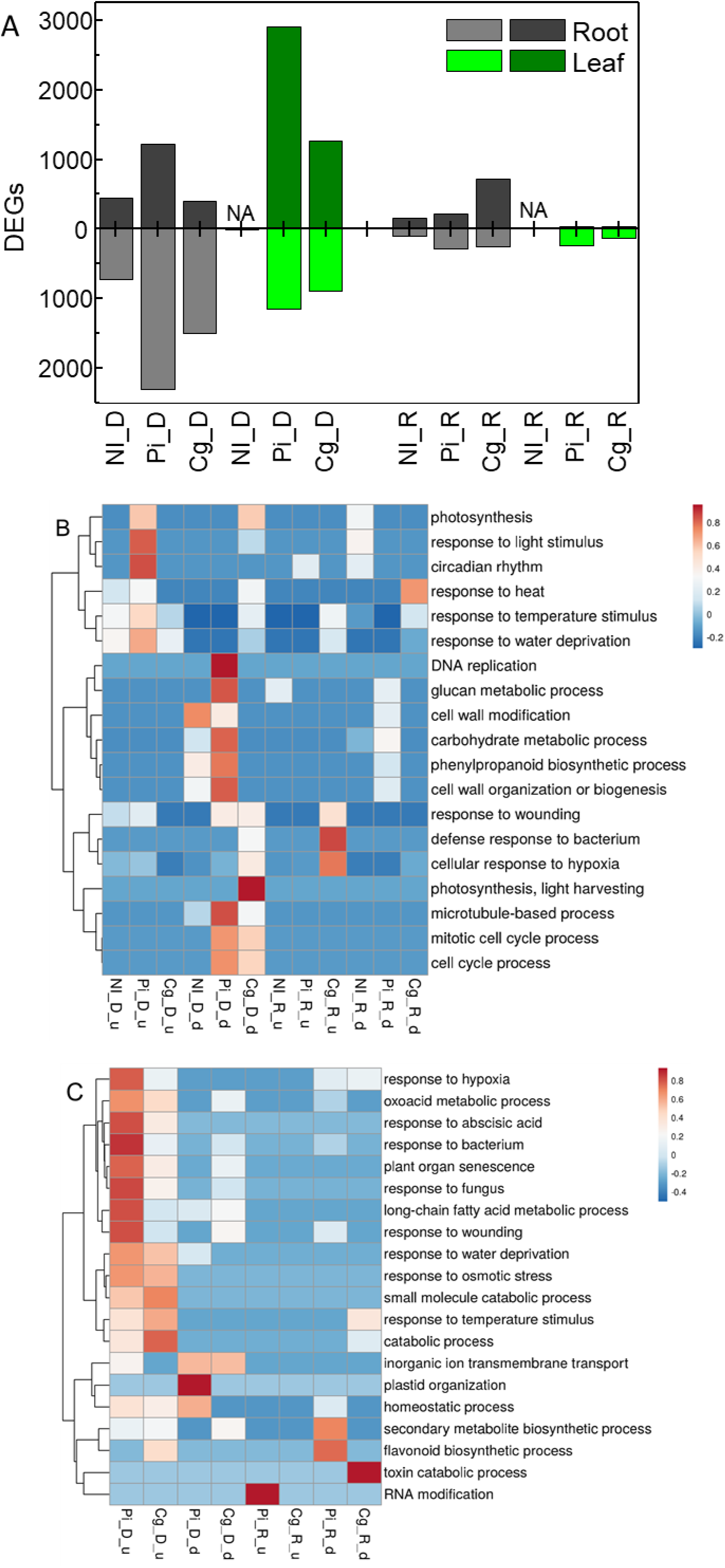
(A) Number of differentially expressed genes (DEGs) in poplars under drought relative to well-watered conditions (D) and in poplars after rewatering relative to well-watered conditions (R). The plants were non-inoculated (Ni) or inoculated either with *P. involutus* (Pi) or with *Cenococcum geophilum* (Cg). NA: not available, number of DEGs ≤ 3. DEGs with Bonferroni adjusted *p* < 0.05 are included. The full list of DEGs is shown in Supplementary Table S2. Light colors indicate down-regulated DEGs and darker colors upregulated DEGs. (B) Selected GO terms in roots and (C) selected GO terms in leaves. Significance of selected GO terms (from Metascape) is indicated by the color for –log10 values. The full list of GO terms for drought responses and recovery is shown in Supplementary Table S3.

To account for differences in the direction of transcriptional changes, we performed separate Gene Ontology (GO) term analyses for up– and down-regulated DEGs (Fig. 4B,C). This approach was necessary because certain GO categories exhibited opposite patterns in Pi and Cg poplars, which would be masked if directionality were ignored. Across drought and recovery treatments in mycorrhizal and non-mycorrhizal plants, numerous enriched GO terms were identified (full lists provided in Supplementary Table S3).

The GO terms underpinned that EM fungal species modulated different types of defense processes. For example,in Pi roots “Response to water deprivation” and “heat” were upregulated and in Cg roots the GO term “Responses to wounding”, “bacteria”, and “hypoxia” were down-regulated (Fig. 4B). These GO terms remained low after recovery (Figure 4B). GO terms for metabolism (carbohydrates, cell wall components, and phenylpropanoids) were suppressed in Pi and Ni roots under drought but remained unaffected in Cg roots (Fig. 4B). In both Pi and Cg roots, cell cycle–related activities were reduced under drought stress. Upon re-watering, most stress-associated GO terms either disappeared or showed markedly reduced significance (Fig. 4B), further supporting a transcriptional shift in roots toward recovery.

Leaves of Ni plants displayed very few DEGs, resulting in no significantly enriched GO terms. In contrast, leaves of Pi and to a lesser extent of Cg plants revealed extensive enrichment of stress-related GO categories under drought (Fig. 4C). Transport processes were down-regulated (Fig. 4C). Following re-watering, stress-related GO terms disappeared, and in Pi roots, “flavonoid biosynthesis” was decreased, whereas “RNA modification” was enriched among upregulated DEGs (Fig. 4C).

### Pi and Cg colonization induce divergent local and systemic transcriptomes in poplar

To identify biological processes showing divergent responses to mycorrhization with Pi or Cg, we directly compared DEGs between the two treatments (Supplementary Tables S4, S5). GO term analysis yielded 126 and 253 enriched categories for leaves and roots, respectively (Supplementary Tables S6, S7).

In leaves under well-watered conditions, no significant GO term was enriched among upregulated DEGs in Pi compared with Cg (Table 3), indicating largely overlapping systemic responses to mycorrhization. Nonetheless, several DEGs were more highly expressed in Pi leaves, including genes for Ca²⁺ signaling, cellulose synthesis, nitrate uptake, and defense responses (e.g., *ERD4*, two potential chaperones, and three putative disease resistance genes) (Supplementary Table S5).

**Table 3:** Significantly enriched GO terms in leaves and roots of Pi and Cg poplars. Significant DEGs for Pi/Cg were used to determine driver GO terms using g_profiler. Data show adjusted p-values for the enrichment. Significant p values are indicated with bold letters.

| **Driver GO terms for leaves** | **Term_id** | Pi_W | Cg_W | PI_D | Cg_D | Pi_R | Cg_R |
| --- | --- | --- | --- | --- | --- | --- | --- |
| response to heat | GO:0009408 | 1 | **4.17E-27** | 1 | **5.63E-23** | 1 | 1 |
| protein folding | GO:0006457 | 1 | **1.86E-25** | 1 | **1.42E-22** | 1 | 1 |
| response to stimulus | GO:0050896 | 1 | **3.31E-14** | **4.16E-29** | **2.52E-14** | 1 | **3.54E-06** |
| response to hypoxia | GO:0001666 | 1 | **1.41E-09** | **6.04E-12** | **2.71E-07** | 1 | 5.19E-01 |
| cellular response to stress | GO:0033554 | 1 | **2.96E-08** | **2.20E-04** | **2.93E-09** | 1 | **7.10E-02** |
| protein phosphorylation | GO:0006468 | 1 | 1 | **1.44E-12** | 1 | 1 | 1 |
| leaf senescence | GO:0010150 | 1 | 1 | **1.41E-05** | 1 | 1 | 1 |
| oxylipin biosynthetic process | GO:0031408 | 1 | 1 | **4.02E-05** | 1 | 1 | 1 |
| jasmonic acid metabolic process | GO:0009694 | 1 | 1 | **1.13E-04** | 1 | 1 | 1 |
| regulation of DNA-templated transcription | GO:0006355 | 1 | 1 | **1.83E-03** | 1 | **6.41E-03** | 1 |
| aromatic compound biosynthetic process | GO:0019438 | 1 | 1 | **3.88E-03** | 1 | **6.19E-02** | 1 |
| secondary metabolite biosynthetic process | GO:0044550 | 1 | 1 | **7.20E-03** | 1 | 1 | 6.41E-01 |
| import across plasma membrane | GO:0098739 | 1 | 1 | **2.95E-02** | 1 | 1 | 1 |
| small molecule biosynthetic process | GO:0044283 | 1 | 1 | **3.30E-02** | 1 | 1 | 1 |
| regulation of nitrogen comp. metabolic process | GO:0051171 | 1 | 1 | 2.52E-01 | 1 | **1.42E-04** | 1 |
| circadian rhythm | GO:0007623 | 1 | 1 | 1 | 1 | **3.67E-02** | **1.18E-03** |
| sulfate assimilation | GO:0000103 | 1 | 1 | 1 | 1 | 1 | **8.14E-04** |
| cellular response to blue light | GO:0071483 | 1 | 1 | 1 | 1 | 1 | **4.46E-03** |
| **Driver GO terms for roots** | **Term_id** | Pi_W | Cg_W | PI_D | Cg_D | Pi_R | Cg_R |
| carbohydrate metabolic process | GO:0005975 | **7.67E-16** | 1 | 1 | **7.46E-11** | 1 | 1 |
| starch catabolic process | GO:0005983 | **5.71E-06** | 1 | 1 | 1 | **1.99E-02** | 1 |
| cell wall organization or biogenesis | GO:0071554 | **1.71E-06** | 1 | 1 | **1.36E-24** | 1 | 1 |
| hemicellulose metabolic process | GO:0010410 | **1.95E-02** | 1 | 1 | **5.23E-12** | 1 | 1 |
| response to stimulus | GO:0050896 | **5.96E-06** | **1.77E-21** | **3.72E-30** | **4.18E-05** | **9.52E-03** | **8.13E-56** |
| response to stress | GO:0006950 | **5.50E-04** | **3.60E-15** | **1.34E-14** | **2.04E-02** | 1 | **2.88E-65** |
| mRNA transcription | GO:0009299 | **1.56E-03** | 1 | 1 | 1 | 1 | 1 |
| maltose metabolic process | GO:0000023 | **1.02E-02** | 1 | 1 | 1 | 1 | 1 |
| multicellular organism development | GO:0007275 | 1.65E-01 | 1 | **2.38E-02** | 1 | 1 | 1 |
| protein folding | GO:0006457 | 1 | **5.94E-12** | 1 | **4.18E-05** | 1 | **5.85E-11** |
| response to acid chemical | GO:0001101 | 1 | **4.15E-05** | **3.71E-07** | 1 | **7.79E-03** | **1.48E-06** |
| biological regulation | GO:0065007 | 1 | **4.40E-04** | **2.04E-10** | 1 | **2.72E-03** | **2.22E-03** |
| transmembrane transport | GO:0055085 | 1 | **7.60E-03** | **1.86E-07** | 1 | **2.18E-05** | 1 |
| regulation of DNA-templated transcription | GO:0006355 | 1 | **1.25E-02** | **6.29E-06** | 1 | **1.32E-02** | 1 |
| water transport | GO:0006833 | 1 | **1.26E-02** | 1 | 1 | 1 | 1 |
| cell communication | GO:0007154 | 1 | **4.67E-02** | **2.24E-04** | 1 | 1 | **2.19E-10** |
| organic cyclic compound biosynthetic process | GO:1901362 | 1 | 5.45E-01 | **1.55E-04** | 1 | **2.12E-02** | 1 |
| potassium ion homeostasis | GO:0055075 | 1 | 7.84E-01 | **2.62E-02** | 1 | 1 | 1 |
| photosynthesis | GO:0015979 | 1 | 1 | **4.20E-46** | 1 | 1 | 1 |
| rhythmic process | GO:0048511 | 1 | 1 | **5.96E-09** | 1 | **1.62E-04** | 1 |
| nitrogen cycle metabolic process | GO:0071941 | 1 | 1 | **1.30E-06** | 1 | 1 | 1 |
| inorganic anion transmembrane transport | GO:0098661 | 1 | 1 | **3.65E-03** | 1 | 4.49E-01 | 1 |
| homeostatic process | GO:0042592 | 1 | 1 | **4.59E-03** | 1 | 1 | 1 |
| plant organ senescence | GO:0090693 | 1 | 1 | **1.28E-02** | 1 | 1 | **5.99E-02** |
| carboxylic acid transport | GO:0046942 | 1 | 1 | **1.80E-02** | 1 | 1 | 1 |
| dipeptide transmembrane transport | GO:0035442 | 1 | 1 | **2.62E-02** | 1 | 1 | 1 |
| intracellular monoatomic cation homeostasis | GO:0030003 | 1 | 1 | **2.68E-02** | 1 | 1 | 1 |
| small molecule metabolic process | GO:0044281 | 1 | 1 | **3.24E-02** | 3.42E-01 | 1 | 3.03E-01 |
| oxoacid metabolic process | GO:0043436 | 1 | 1 | 1.07E-01 | 1 | 1 | **1.54E-02** |
| phenylpropanoid metabolic process | GO:0009698 | 1 | 1 | 1 | **6.17E-10** | 1 | 1 |
| protein phosphorylation | GO:0006468 | 1 | 1 | 1 | **4.16E-03** | 1 | **9.83E-04** |
| camalexin biosynthetic process | GO:0010120 | 1 | 1 | 1 | 1 | 1 | **1.28E-02** |
| organonitrogen compound metabolic process | GO:1901564 | 1 | 1 | 1 | 1 | 1 | **1.35E-02** |

Under drought, Pi leaves showed enrichment for stress-related GO terms (“response to stimulus,” “response to hypoxia,” “cellular response to stress”), alongside “senescence,” “transport stimulation,” “jasmonate metabolism,” and “secondary compound metabolism” (Table 3). In contrast, Cg leaves already displayed stress-related GO terms under well-watered conditions, together with “response to heat” and “protein folding” (Table 3), indicating distinct stress-priming compared with Pi plants. Among the most strongly regulated Cg genes was *GALACTINOL SYNTHASE* (*GolS*), known for its role in drought protection (Sengupta et al., 2015). Stress-associated GO terms in Cg leaves remained stable under drought. After re-watering, most stress terms disappeared, while categories related to circadian rhythm, nitrogen metabolism, and sulfur metabolism emerged (Table 3).

Root responses differed markedly from leaves. GO terms “response to stimulus” and “response to stress” were enriched across nearly all conditions (Table 3). “Protein folding” was uniquely enriched in Cg roots across all treatments, representing a major distinction from Pi roots. In well-watered Pi roots, carbohydrate metabolism and cell wall–related GO terms were enriched but disappeared under drought, whereas these processes appeared in stressed Cg roots. Well-watered Cg roots showed enrichment for cell communication and water transport, which under drought were lost but appeared in Pi roots.

Additional drought-responsive GO terms in Pi roots included cation/anion transport, senescence, and other cellular processes, while stressed Cg roots showed enrichment for “phenylpropanoid metabolism” and “protein phosphorylation.” After re-watering, only a few broad GO terms (e.g., “response to acid chemical,” “regulation of biological processes”) remained in both mycorrhizal types. However, Cg roots retained strong enrichment for “response to stimulus,” “response to stress,” and “protein folding,” further supporting a persistent stress-priming effect.

### Heat shock proteins and aquaporins support stress preparedness of Cg poplars

Since “protein folding” was the most distinctive GO term in Cg compared with Pi poplars, we examined the transcriptional profiles of DEGs in this category for each condition (Fig. 5). Hierarchical clustering revealed two major groups: (1) chaperonin-like genes, including putative calreticulins and *ERD10*, generally more highly expressed in leaves than in roots; and (2) heat shock proteins (*HSP*s), cyclophilin-like genes, and *DNAJ*, with transcript levels typically higher in roots than in leaves (Fig. 5). In Cg plants, these genes showed consistently higher transcript abundances than in Ni or Pi plants under both well-watered and drought conditions. Network analysis indicated that “protein folding” DEGs formed a highly co-expressed cluster (45 nodes, 429 edges; expected edges = 25; average node degree = 19.1; clustering coefficient = 0.7; p < 10⁻¹⁶; Supplementary Fig. S3). Since the enrichment of GO term for “heat stress” and “protein folding” was surprising, we confirmed these responses to Cg mycorrhization in the absence of stress in poplar plantlets grown under strictly temperature-controlled conditions (24°C) in sterile Petri dish systems (Supplement Table S8).

**Figure 5.**
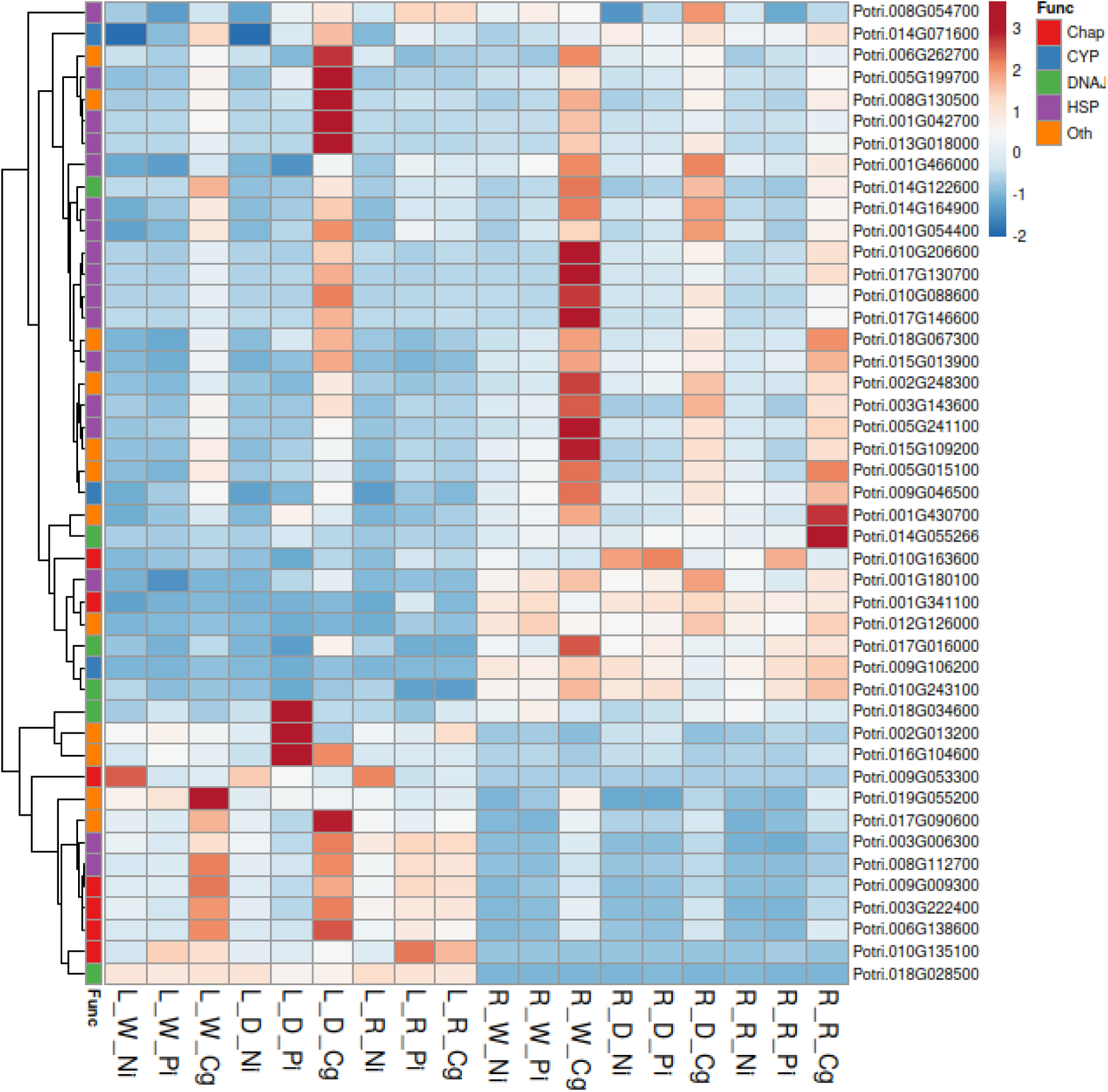
Hierarchical clustering of mean transcript abundances of genes putatively involved in protein folding and heat shock responses. A total of 45 DEGs were identified in the data set and their transcript abundances are shown across all conditions: L = leaves, R: Roots, _W = well-watered, _D = drought, _R = rewatered,_NI = non-inoculated, Pi = P*. involutus*, C = *C. geophilum*. The first column shows a classification of putative gene functions: CHAP = chaperone/chaperonin-like, HSP = heat shock protein, CYP = cyclophilin-like protein, DNAJ = co-chaperones, Oth = other functions.

Given physiological evidence for differences in water potential and GO enrichment in “transport processes” and “water transport” in Cg versus Pi poplars, we further analyzed transcriptional profiles of aquaporin DEGs (Fig. 6). Putative aquaporins were annotated as plasma membrane intrinsic proteins (PIPs), tonoplast intrinsic proteins (TIPs), nodulin-26 intrinsic proteins (NIPs), and small basic intrinsic proteins (SIPs) (Gupta and Sankararamakrishnan, 2009). Expression patterns varied strongly with stress treatment, EM species, and tissue, but most aquaporins were more highly expressed in roots than in leaves (Fig. 6). Notably, Cg roots showed markedly higher expression of multiple PIPs and TIPs compared with Pi or Ni roots under well-watered and recovery conditions (Fig. 6).

**Figure 6.**
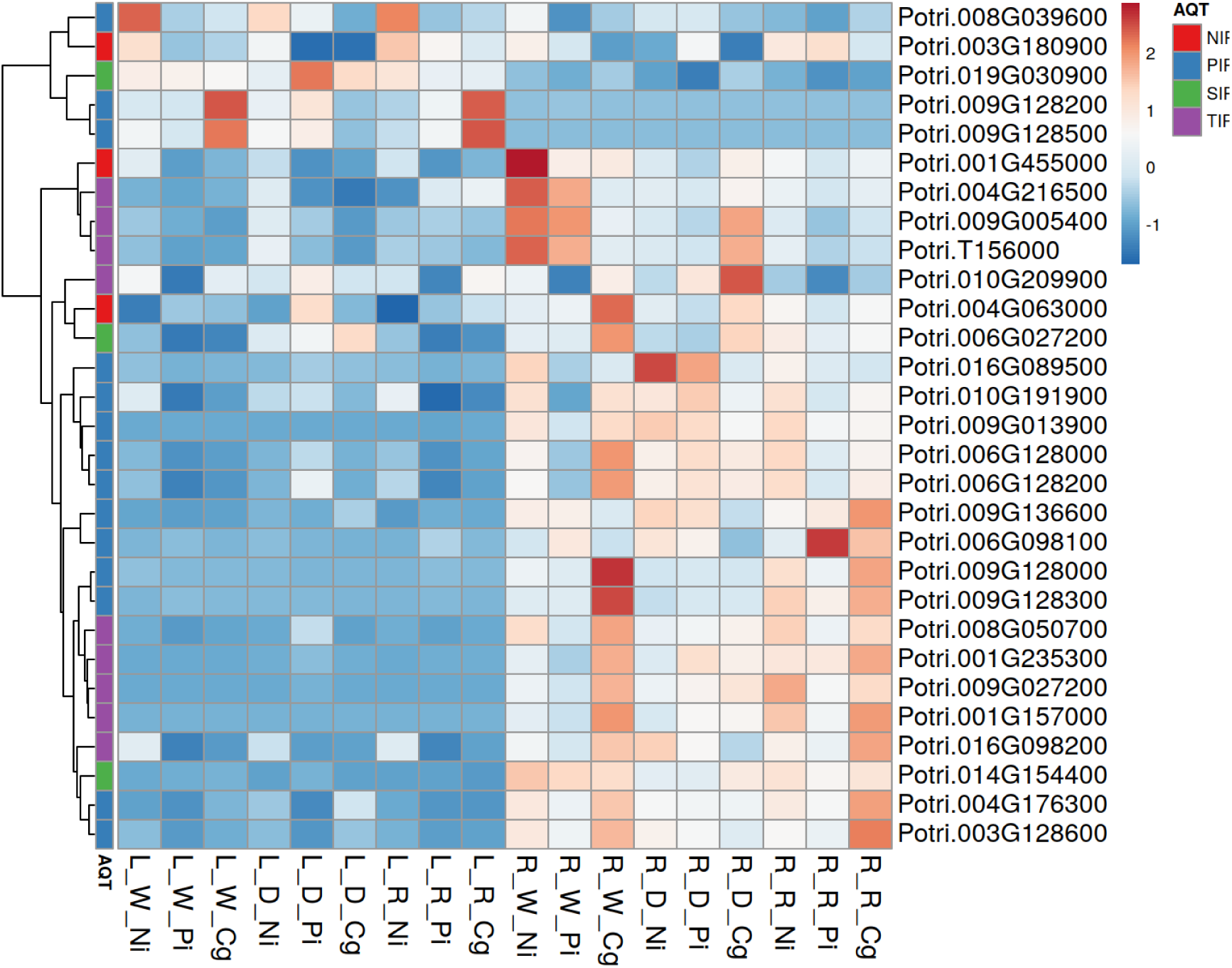
Hierarchical clustering of mean transcript abundances of putative aquaporin-like genes. A total of 29 DEGs were identified in the data set and their transcript abundances are shown across all conditions: L = leaves, R: Roots, _W = well-watered, _D = drought, _R = rewatered,_NI = non-inoculated, Pi = P*. involutus*, Cg = *C. geophilum*. The first column shows a classification of the putative aquporin types (AQT): NIP = nodulin-26 intrinsic protein, SIP = small basic intrinsic protein, PIP = plasma membrane intrinsic protein, TIP = tonoplast intrinsic protein

### Growth regulation and leaf shedding contribute to poplar drought acclimation

In Arabidopsis, growth-defense trade-off is regulated by the TOR/RAPTOR (TARGET OF RAPTAMYCIN/REGULATORY-ASSOCIATED PROTEIN OF TOR) complex (growth stimulation) and its negative regulator SnRK1 (Sucrose non-fermenting-1-related protein kinase) (Margalha et al., 2019; Jamsheer K et al., 2022). Fine tuning occurs via the FLZ8 protein (Jamsheer K et al., 2022). We identified poplar homologs of these genes (Fig. 7A) and found that putative TOR/RAPTOR transcripts in leaves were suppressed under drought in both Pi and Cg plants, and were also low in Cg under well-watered conditions. Putative *SnRK1* transcripts peaked under drought in Cg leaves, while FLZ8 expression increased in both mycorrhizal types under drought. Assuming similar functions as in Arabidopsis, *TOR/RAPTOR* regulation is consistent with higher leaf growth in Pi compared with Cg under well-watered conditions, and suppression of Pi leaf growth under drought.

**Figure 7:**
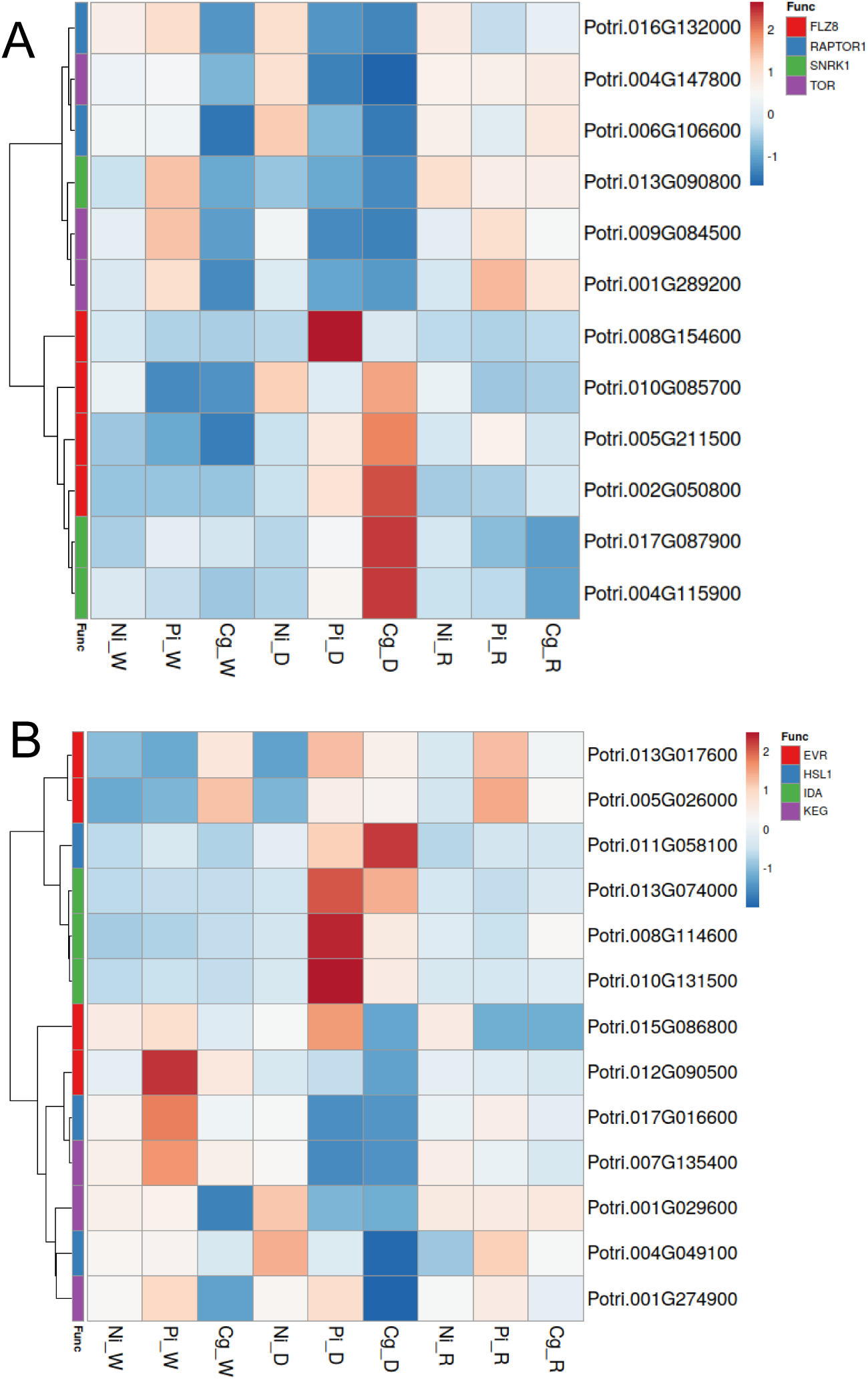
Hierarchical clustering of mean transcript abundances of genes potentially involved in growth regulation (A) and leaf shedding (B). Mean transcript abundances are shown across all conditions in leaves. Pi = P*. involutus*, Cg = *C. geophilum,* _W = well-watered, _D = drought, _R = rewatered, _NI = non-inoculated. The first column shows a classification of putative functions (Func): FLZ = Fine tuning protein, SnRk1 = SNF Kinase, TOR = TARGET OF RAPAMYCIN, RAPTOR = REGULATORY ASSOCIATED PROTEIN OF TOR, EVR = EVERSHED, HSL = receptor-like kinase, IDA = INFLORESCENCE DEFICIENT ABSCISSION, KEG = KEEP on GOING.

Under water scarcity, poplars often limit water consumption through controlled leaf abscission (Fischer and Polle, 2010), involving signaling via receptor-like protein kinases (HSL [HAE] and EVERSHED [EVR]), MKKs (MITOGEN-ACTIVATED PROTEIN KINASEs) including KEGs (KEEP ON GOING) and IDA (INFLORESCENCE DEFICIENT ABSCISSION) (Patharkar and Walker, 2018). In our study, all detected transcripts for *IDA* homologs were significantly upregulated under drought in Pi compared with Cg plants (Fig. 7B). Conversely, one potential one potential *HSL1* gene (Potri.011G058100), whose homolog is positively associated with longevity in Arabidopsis (Chen et al., 2022), showed higher transcript abundances in Cg than in Pi (Fig. 7B).

## Discussion

### *P. involutus* mycorrhizas benefit poplar growth but not drought resistance

Regulation of growth and stress acclimation is critical for plant survival in dynamic environments. A key finding of our study was that colonization by different EM fungi triggered sharply contrasting local and systemic transcriptional responses, with divergent consequences for host growth and defense under both non-stressed and stressed conditions. This highlights the remarkable plasticity of poplar to adjust to environmental challenges.

Previous work has shown that growth responses to mycorrhizal colonization vary greatly depending on the specific plant–fungus combination (Shinde et al., 2018; Szuba et al., 2019; Dagher et al., 2020; Birch et al., 2021; Da Costa et al., 2022; Shi et al., 2024). Growth stimulation by EM fungi is often linked to enhanced nutrient uptake, particularly nitrogen (Pena and Tibbett, 2024). Consistent with earlier studies (Luo et al., 2009; Marqués-Gálvez et al., 2025), we observed increased transcription of transporters for nitrate, ammonium, and amino acids in Pi-colonized poplars, supporting elevated N metabolism. However, tissue N concentrations were lower in Pi plants than in those colonized by Cg. Since photosynthetic carbon assimilation did not differ between Pi and Cg poplars, and Cg plants had higher tissue N but lower growth, these results indicate that Pi increased nitrogen use efficiency, allocating available carbon more effectively to growth over other sinks.

Pi-induced growth stimulation was accompanied by activation of growth regulators, including the TOR system, cellulose biosynthesis, and phytohormone metabolism, all known to interact during leaf development (Xiong et al., 2021). Leaf area, the dominant determinant of biomass production (Pellis et al., 2004; Yu et al., 2019), was strongly increased in Pi poplars. Thus, distinct transcriptional rewiring appears central to the growth enhancement of Pi compared with Ni and Cg plants.

EM colonization also reprograms plant defense, elevating proteins and metabolites such as protease inhibitors, chitinases, aldoximes, and phenolic compounds (Kaling et al., 2018; Sebastiana et al., 2018; de Freitas Pereira et al., 2023; Marqués-Gálvez et al., 2025; Srivastava et al., 2025). While such systemic responses enhance resistance to biotic stressors (Kaling et al., 2018; Vishwanathan et al., 2020), their role in mitigating abiotic stress is less clear. Under moderate drought, physiological and transcriptional responses did not differ between non-mycorrhizal and *L. bicolor*-colonized poplars (de Freitas Pereira et al., 2023). Under severe drought in this study, Pi and Ni plants exhibited similar declines in stomatal conductance and photosynthesis, but Pi poplars experienced more severe reductions in leaf water potential and biomass. Leaf shedding is a common drought acclimation strategy that reduces water loss (Wolfe et al., 2016). In line with greater leaf loss, Pi plants showed significant upregulation of *IDA* homologs involved in controlling abscission (Patharkar and Walker, 2018). The enhanced leaf area of Pi plants likely required tighter regulation of canopy size to balance water demand with declining soil moisture. This may explain drought-induced transcriptomic differences between Ni and Pi plants: Pi poplars engaged a more pronounced stress acclimation program to survive. Importantly, Pi plants resumed post-stress growth more rapidly than Ni plants, suggesting a potential long-term ecological advantage despite lower drought resistance.

### *C. geophilum* mediates drought tolerance at the expense of growth

Our results show that Cg colonization caused profound physiological and molecular changes in poplar even under optimal water supply. Unlike Pi and Ni plants, Cg-colonized poplars exhibited clear growth reduction in the absence of drought. This was not linked to impaired photosynthetic carbon assimilation or nitrogen uptake but rather to transcriptional evidence of a resource shift from growth to defense and stress readiness under well-watered conditions.

A key mechanism supporting this shift was strong upregulation of *GolS*, an enzyme in raffinose family oligosaccharide biosynthesis (Sengupta et al., 2015). *GolS* promotes the accumulation of osmolytes such as galactinol, myo-inositol, and raffinose, which aid osmotic adjustment and abiotic stress protection (Unda et al., 2017; La Mantia et al., 2018; Liu et al., 2021; Shikakura et al., 2022). Overexpression in poplar enhances stress tolerance but reduces growth (Unda et al., 2017), a trade-off also evident in Cg plants. This suggests strategic carbohydrate allocation to stress-protective compounds rather than biomass production.

Cg poplars also showed massive aquaporin upregulation, particularly in roots. Aquaporins, membrane channels that facilitate water movement, are critical for maintaining turgor under water deficit and for efficient root water uptake in both plants and fungal symbionts (Secchi and Zwieniecki, 2013; Xu et al., 2015; Xu et al., 2016; Jiang et al., 2020; Tomkins et al., 2021). The high aquaporin expression in Cg tissues (Peter et al., 2016; our study), likely enhanced root water permeability and helped maintain leaf water potentials during drought.

A notable, unexpected finding was strong induction of multiple heat shock transcription factors (*Hsfs*) and heat shock proteins (*HSP*s) in Cg roots and leaves, a response absent in Pi and Ni plants. HSPs act as molecular chaperones, protecting protein integrity during heat and other abiotic stresses (Jacob et al., 2017; Tian et al., 2021), but their ATP-dependent activity is energetically costly (Berka et al., 2022). *GolS* is directly regulated by HsfA2 (Nishizawa et al., 2006) and genetic studies show that enhanced *HsfA2* expression induces *GolS* and improves tolerance to high light, oxidative stress, and heat (Nishizawa-Yokoi et al., 2009; Song et al., 2016; Gu et al., 2019). Our results support a coordinated defense mechanism between Hsfs and GolS in Cg plants that is active even without abiotic stress.

Collectively, Cg colonization was associated with suppression of cell cycle activity and the *TOR/RAPTOR* growth-regulatory module (Liu and Xiong, 2022), which in Arabidopsis is glucose-regulated and influences *HSP* expression (Sharma et al., 2019). These changes point to transcriptional and cellular resource reallocation that prioritizes stress preparedness over growth.

### Conclusions – ecological and mechanistic implications

Our findings reveal two contrasting mycorrhiza-mediated drought strategies in poplar. Pi promotes vigorous growth under optimal conditions, combined with rapid drought-induced leaf abscission that reduces water loss and enables swift recovery. In contrast,Cg induces a constitutive “pre-armed”state, characterized by upregulation of osmolyte biosynthesis, aquaporins, and Hsf/HSP chaperone networks. This strategy safeguards water status during drought but consistently limits growth. These contrasting approaches exemplify the growth–defense trade-off theory (Huot et al., 2014; He et al., 2022), demonstrating that EM fungi occupy distinct positions along this spectrum. Our results connect ecological traits with molecular evidence (Monson et al., 2022) and highlight the potential of tailoring mycorrhizal partnerships to specific environmental challenges: selecting partners like Pi to maximize biomass in favorable climates, or Cg to ensure resilience under chronic water limitation. Such species-specific symbiotic engineering offers a promising avenue for next-generation forestry and crop breeding in the face of a changing climate.

## Materials and methods

### Cultivation of poplar and ectomycorrhizal fungi under sterile conditions

*Populus × canescens* (a hybrid of *Populus tremula* x *Populus alba*, INRA 717 1B4) plantlets were multiplied by *in vitro* micropropagation and grown on half-strength MS medium (Murashige and Skoog, 1962) in glass jars for approximately three weeks under controlled environmental conditions [air temperature: 24 °C, 16h/8h light/dark cycle with 75 μE m^−2^ s^−1^ photosynthetic active radiation (PAR)]. Two weeks after poplar propagation, fungal cultures of *Paxillus involutus* (Batsch., strain MAJ) and *Cenococcum geophilum* (Fr., strain DR 041) were started. Per fungal culture, 40 ml of solid modified Melin–Norkrans (MMN) medium (0.34 mM CaCl_2_, 0.43 mM NaCl, 3.67 mM KH_2_PO_4_, 1.89 mM (NH_4_)_2_SO_4_, 0.61 mM MgSO_4_·7H_2_O, 0.037 μM FeCl_3_·6H_2_O, 0.3 μM thiamine HCl, 1.1% glucose, 0.3% malt extract (Merck, Darmstadt, Germany), 1% agar (Duchefa, Haarlem, Netherlands), pH 5.2) was prepared in a Petri dish (12 × 12 cm). In each Petri dish, two sterilized cellophane membranes (6 × 12 cm, Deti GmbH, Meckesheim, Germany) were placed on the solid medium. Then, five fungal plugs (each approximately 0.25 cm^2^) were distributed on each cellophane membrane and cultivated for approximately 10 days at 20 °C in darkness. Controls were prepared with cellophane membranes on MMN medium without fungal inoculation. For the co-cultivation of poplar with different ectomycorrhizal fungal isolates, we used the sandwich system described by Müller et al., (2013): the rooted plantlets were transferred to Petri dishes, which were half-filled with solid medium (40 ml modified M-MMN-S mixed medium: 0.34 mM CaCl_2_, 0.43 mM NaCl, 3.67 mM KH_2_PO_4_, 1.89 mM (NH_4_)_2_SO_4_, 0.61 mM MgSO_4_·7H_2_O, 0.037 μM FeCl_3_·6H_2_O, 0.3 μM thiamine HCl, 0.67 mM MnSO_4_·H_2_O, 0.76 mM ZnSO_4_·7H_2_O, 0.20 mM CuSO_4_·5H_2_O, 18.61 μM (NH_4_)_6_Mo_7_O_24_·4H_2_O, 0.2% sucrose, 2% agar, pH 5.5) covered with a sterilized cellophane membrane. The whole plantlet was placed inside the Petri dish with the roots on the membrane. The roots were then covered with a prepared cellophane membrane, either with or without fungal mycelium. The mycelium was in contact with the roots. After sealing with gas-permeable Parafilm^TM^ (Bemis Flexible Packing, Neenah WI, USA), the Petri dishes were positioned vertically in racks and the lower halves were wrapped in aluminum foil to keep poplar roots and fungi in darkness. The co-culture systems with poplar and fungi were incubated in environmental cabinets (Percival, Emersacker, Germany) for three weeks at 23 °C with a 16h/8h light/dark cycle, 80 μE m^−2^s^−1^ PAR and 60% relative air humidity. During the co-culture with and without different EM fungi, poplars in the Petri dishes were inspected regularly and contaminated plants were immediately disposed. Roots of inoculated spare plants were inspected for the formation of EM structures under a microscope (DFC 420, Leica Microsystems, Wetzlar, Germany). Pi-colonized root tips were ensheathed by silvery-shining and Cg-colonized by black hyphae, which extended from the fungal plugs. EM roots were shorter, had a rounded root tip and less root hairs than Ni roots. After three weeks, the plants were well colonized.

In an independent experiment, poplar plants were either co-cultivated with *C. geophilum* (three plugs) or grown without inoculation in sterile plate systems as described above. After 20 days, the leaves of the sterile-grown plantlets were harvested, flash-frozen in liquid nitrogen and stored at –80°C.

### Poplar cultivation and drought treatment

Colonized and Ni poplar plants were potted individually into 3L pots containing a mixture of coarse sand (Ø 0.71-1.25 mm), fine sand (Ø 0.4-0.8 mm, Dorfner GmbH, Hirschau, Germany) and peat (Fruhstorfer Erde ‘Nullerde’, Hawita Gruppe GmbH, Vechta, Germany) with the ratio of 8:2:2 v/v. Before use, the sand was washed, then mixed with peat and the mixture was autoclaved (HST 6 × 6 × 6, Zirbus Technology GmbH, Bad Grund, Germany) twice at 120°C for 20 minutes. Before potting, five additional 2-week-old fungal plugs were added to the soil of the mycorrhizal poplars. The plants exhibited mean heights of 6.2 ± 0.4 cm, irrespective of EM or Ni treatments. To acclimate the plants to greenhouse conditions, they were covered with transparent beakers, which were gradually lifted and removed after 2 weeks. Then, plants were grown under semi-controlled greenhouse conditions at 50% relative air humidity and air temperatures within a range of lower and upper thresholds of 18°C and 28°C. Some outlier days with up 35°C occurred, when bright sun irradiation (no clouds) lasted more 12h in summer. The plants were rotated regularly to avoid positional effects and distributed across three greenhouse cabinets. Each cabinet contained plants of each inoculation treatment (Pi, Cg and Ni poplars). Ambient light was supplemented for 16h per day with 180 μE m^−2^s^−1^ PAR (L18W/840, Osram, Munich, Germany). The plants were irrigated daily with 200 ml distilled water and 50 ml modified Long Ashton nutrient solution (500 μM KNO_3_, 900 μM Ca (NO_3_)_2_, 300.2 μM MgSO_4_, 59.99 μM KH_2_PO_4_, 4.13 μM K_2_HPO_4_, 10 μM H_3_BO_3_, 2 μM MnSO_4_, 7 μM Na_2_MoO_4_, 50 nM CoSO_4_·7H_2_O, 200 nM ZnSO_4_·7H_2_O, 200 nM CuSO_4_·5H_2_O, 10 μM ethylenediamine tetraacetic acid (EDTA)-Fe, pH 5.8). After 16 weeks under these conditions, each group (Pi, Cg and Ni poplars) was divided into 3 subgroups: well-watered, drought-exposed, and re-watered plants. The total number of replicates was 6 per fungal treatment and subgroup. The experimental period lasted 4 weeks: the well-watered plants were irrigated as before (daily 200 ml water, 50 ml nutrient solution); drought-treated plants received daily 50 ml of nutrient solution and a subset of these plants was re-watered for one week before harvest. All plants were harvested 20 weeks after potting.

### Growth and physiological measurements

During cultivation in the greenhouse, plant height was recorded once a week, and stem diameter and leaf numbers twice a week. The absolute growth rates (AGR) of plant height and stem diameter were calculated with the following equation:

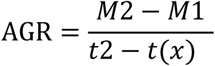

M1 and M2 are plant height (cm) or stem diameter (mm) at the time points, t_x_ and t2, respectively with t_(x1)_ = 14d for well-watered plants (starting when the plants were acclimated to the ambient conditions), t_(x2)_ = 118d for the drought-stressed plants (encompassing the time of low soil water contents) and t_(x3)_ = 132d (time-point when rewatering started). For all treatments t2 = 139d, i.e., the day of harvest.

During the experimental period, the soil moisture of each pot was monitored daily with a tensiometer (HH2 Moisture Meter version 2.3, Delta-T Devices, Cambridge, UK) at a depth of 10 cm.

Gas exchange (net photosynthesis, transpiration, and stomatal conductance) was measured weekly on the first fully light-exposed developed leaf (5^th^ leaf counted from the top) with a portable photosynthesis system (LI-6800, LI-COR Biosciences GmbH, Bad Homburg, Germany).

The pre-dawn leaf water potential was measured on the fully developed leaves at the top (approximately leaf number 6) using a Scholander pressure chamber (M 1505D, PMS instrument, Albany, USA) from 3 am to 5 am, before sunrise.

Shed leaves from each plant were counted, collected and dried (60°C for 7 days) to determine biomass loss.

### Harvest and biomass determination

On the day of the harvest, three leaves were collected (one from the top, the middle, and bottom of the stem) of each plant, weighed fresh, scanned, dried and weighed again. The scans were used for leaf area determination with Image J (https://imagej.net/ImageJ). Average leaf size and whole-plant leaf area of each plant were calculated with the following equations (Yu et al., 2021):

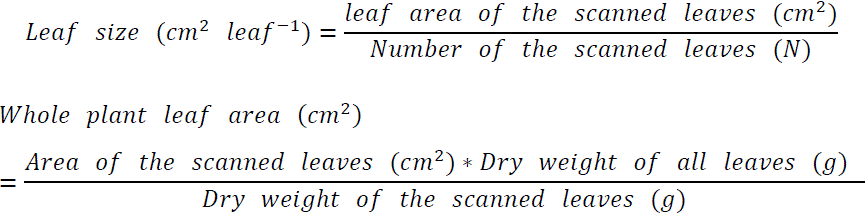

Approximately eight leaves (from the top) and a 3-cm stem section from the bottom were weighed, wrapped in aluminum foil bags, immediately frozen in liquid nitrogen, and stored at −80 °C. The remaining leaves and stem were weighed, dried at 60 °C for two weeks, and weighed again. The roots with the attached soil were carefully removed from the pot and immersed briefly in water to gently wash-off sand and peat. Then, the roots were quickly separated into fine (< 2 mm) and coarse roots (> 2 mm), surface-dried between tissue paper, and weighed. One part of the fine roots was shock-frozen in liquid nitrogen and stored at −80 °C. Three fine root segments were collected randomly from each plant and transferred into FAE (37% formalin, 100% glacial acetic acid, 70% ethyl alcohol = 5:5:90) for microscopic investigations (Luo et al., 2009). The remaining root fractions were dried at 60 °C for 2 weeks and used for determination of dry weight.

The fresh and dry mass of a tissue was used to determine the dry-to-fresh weight ratio. The fresh weights of all aliquots per tissue were added and used to determine the dry biomass of a tissue:

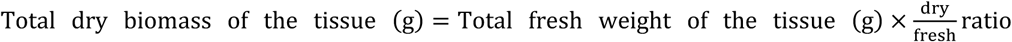

Total plant dry biomass (g) was the sum of leaf, stem, coarse and fine root dry biomass.

### Determination of ectomycorrhizal colonization of the roots

Fine root samples, preserved in FAE, were spread individually in Petri dishes with distilled water and observed under a microscope (Stemi SV11, Zeiss, Jena, Germany). All mycorrhizal and non-mycorrhizal root tips per sample were counted and EM colonization was calculated for each plant:

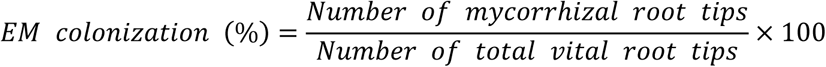

### RNA extraction and sequencing

Leaves and roots of three biological replicates per treatment were used for RNA isolation. Briefly, about 300 mg of frozen sample was milled to a fine powder in ball mill (MM400, Retsch, Haan, Germany) and used for RNA extraction with the CTAB (hexadecyltrimethylammonium bromide) method (Chang et al., 1993). The TURBO DNA-free^TM^ kit (Thermo Fisher Scientific, Waltham, USA) was used to purify the RNA.

The concentration of purified RNA from leaves and roots ranged from 250 to 330 ng µL^-1^ and was diluted to ratio of A260/A280 of 2.0, which was the requested standard for RNA sequencing. Library preparation and RNA sequencing were performed at the NGS-Integrative Genomics Core Unit (NIG), Department of Human Genetics, University Medical Center Göttingen (UMG). At least 800 ng RNA of each sample was used for library preparation with the TruSeq mRNA Sample Prep kit v2 (Illumina, San Diego, CA, USA) and 50 bp single-end sequences were generated on a HiSeq4000 sequencer (Illumina, San Diego, CA, USA).

### Bioinformatic analysis of RNAseq Data

Raw reads were filtered and trimmed with Fastp version 0.21.0 using the default settings (Chen et al., 2018). After processing, approximately 20 to 45 million reads remained per sample. The processed reads from each library were aligned to the reference genome of *Populus trichocarpa* v3.1 (Tuskan et al., 2006) using HISAT2 version 2.1.0. Raw gene counts were generated with FeatureCounts version 2.0.0. Normalization of gene counts per sample and identification of differentially expressed genes (DEGs) between the treatments were calculated with the R package DESeq2 version 1.32.0 (Love et al., 2014). We used DEGs with log2-fold change > ǀ1.0ǀ and Bonferroni adjusted *p* values < 0.05 for further analyses of the greenhouse experiment and all DEGs for the sterile grown plantlets. Gene ontology (GO) enrichment analyses for the category “Biological Processes” were conducted with g:Profiler (https://biit.cs.ut.ee/gprofiler/gost), running multiquery analyses (Kolberg et al., 2023). In addition to complete lists, the analyses calculates “driver” GO terms, which represent significant categories of similar GO terms. We also analysed GO terms with Metascape (https://metascape.org/; (Zhou et al., 2019)) and used the proposed list of selected GO terms. The best AGI matches of the Potri IDs were used as input data (Supplemental Table S1). To provide an overview, we used GO terms (“driver” GO obtained by “gprofiler” analysis), which represent significant categories of similar GO terms. Heatmaps were generated with mean counts with Clustvis (https://biit.cs.ut.ee/clustvis/) (Metsalu and Vilo, 2015) applying the following settings: rows are centered; unit variance scaling is applied to rows. Rows are clustered using correlation distance and average linkage.

### Carbon and nitrogen analyses

Dry plant tissue including leaves, stem, coarse and fine roots were ground to a homogenous fine powder in a ball mill (MM400, Retsch, Haan, Germany). Approximately 2 mg of leaves, stem and fine roots, 3 mg of coarse roots were weighed into 4 mm x 6 mm tin cartouches (IVA Analysentechnik, Meerbusch, Germany) using a super-micro balance (S4, Sartorius, Goettingen, Germany). Carbon (C) and nitrogen (N) contents were measured with an element analyzer (Flash EA 1112, Thermo-Electron, Milano, Italy). Acetanilide (Sigma-Aldrich) was used as the standard.

The N concentration (N) (mg g^-1^) and the biomass of each tissue (g) were used to determine the whole-plant N content as

Whole-plant N content (mg plant^-1^) = N_leaf_ x biomass_leaf_ + N_stem_ x biomass_stem_ + N_coarse root_ x biomass_coarse root_ + N_fine root_ x biomass_fine root_

The weighted mean N concentration was determined as

Weighted mean N concentration (mg g^-1^ biomass) = Whole-plant N content (mg) / whole plant dry biomass (g)

To determine nitrogen use efficiency for the whole growth period in the greenhouse (139 days), we plotted whole-plant biomass against whole-plant N content and determined the slopes of linear models for Ni, Pi and Cg. Differences between the regression lines were analyzed with the function “comparison of regression lines” of the program Statgraphics Centurion (version 18.1.12, Statgraphics Technologies Inc., The Plains, Virginia, USA).

### Statistical Analysis

Data are shown as means (n = 5 or 6 biological replicates, ± SE). Differences between means were tested with software program R version 4.0.2 (R DEVELOPMENT CORE, 2020). Normal distribution and homogeneity of the variances were assessed visually using histograms and plotting the residuals. We considered *p*-values < 0.05 obtained by two-way ANOVA and post hoc Fisher’s LSD test to indicate significant differences between the means per treatment. Percentage data such as EM colonization rates were processed with a beta regression model and then used to detect significant differences at *p* < 0.05 with ‘Tukey adjusted comparison’.

## Acknowledgements

We are grateful to Merle Fastenrath, Gabriele Lehmann and Cathrin Leibecke for excellent technical assistance with the plant and fungal cultivation, to Lars Szwec (Centre for Stable Isotope Research and Analysis, University of Göttingen) for the C and N analyses, and to Thomas Klein (Laboratory for Radioisotopes) for skillful RNA extraction. We also thank the NGS-Integrative Genomics Core Unit (NIG), Department of Human Genetics, University Medical Center Göttingen (UMG) for their help with the library preparation and sequencing and to Dr. Johannes Ballauff and Jacob Schmidt for processing RNAseq data.

## Author contributions

HS conducted the experiments, analyzed physiological and bioinformatics data, wrote the first draft, ZL conducted the plate experiments and bioinformatic analyses, AP developed the concept, analyzed data, wrote the final draft, and secured funding. All authors revised the paper and agreed on its final version.

## Data availability

RNA-seq raw data have been deposited in the ArrayExpress database at EMBL-EBI (www.ebi.ac.uk/arrayexpress) under accession number E-MTAB-12863. The data tables with annotations and differential analyses for all treatment combinations is available (after acceptance) in Figshare under 10.6084/m9.figshare.30132064.

## Supplementary materials

**Supplementary Figure S1:** Diameter growth rate of mycorrhizal poplars in response to well-watered, drought and rewatered treatments.

**Supplementary Figure S2:** Relationship between poplar N content and biomass.

**Supplementary Figure S3**: Network analysis of DEGs in the GO term „Protein Folding“.

**Supplementary Table S1:** Leaf number, size, and tissue biomass of poplar after mycorrhization with *P. involutus* (Pi), *C. geophilum* (Cg) or non-inoculated plants (Ni) in response to well-watered, drought-stressed and rewatered conditions.

**Supplementary Table S2**: Log2-fold changes of transcript abundances and Bonferroni adjusted p values for the comparisons among fungal treatments and well-watered, drought-stressed and rewatered conditions.

**Supplementary Table S3:** All GO terms for up– and down-regulated DEGs in roots and leaves of drought-stressed and rewatered poplar plants relative to well-watered plants.

**Supplementary Table S4:** Differentially expressed genes *Paxillus involutus-* relative to *Cenococcum geophilum*-colonized roots.

**Supplementary Table S5:** Differentially expressed genes in leaves of *Paxillus involutus-* relative to *Cenococcum geophilum*-colonized plants.

**Supplementary Table S6:** Complete list of significantly enriched GO terms in roots of Cg– and Pi-colonized plants under well-watered, drought-stressed and rewatered conditions.

**Supplementary Table S7:** Complete list of significantly enriched GO terms in leaves of Cg– and Pi-colonized plants under well-watered, drought-stressed and rewatered conditions.

**Supplementary Table S8:** Significant GO term for DEGs in leaves of *P.* x *canescens* grown in sterile plate systems in the presence or absence of *C. geophilum*.

