## Supplemental Figures for "Divergent Strategies of Mycorrhiza-Mediated Drought Adaptation in Poplar"

### Supplementary figures

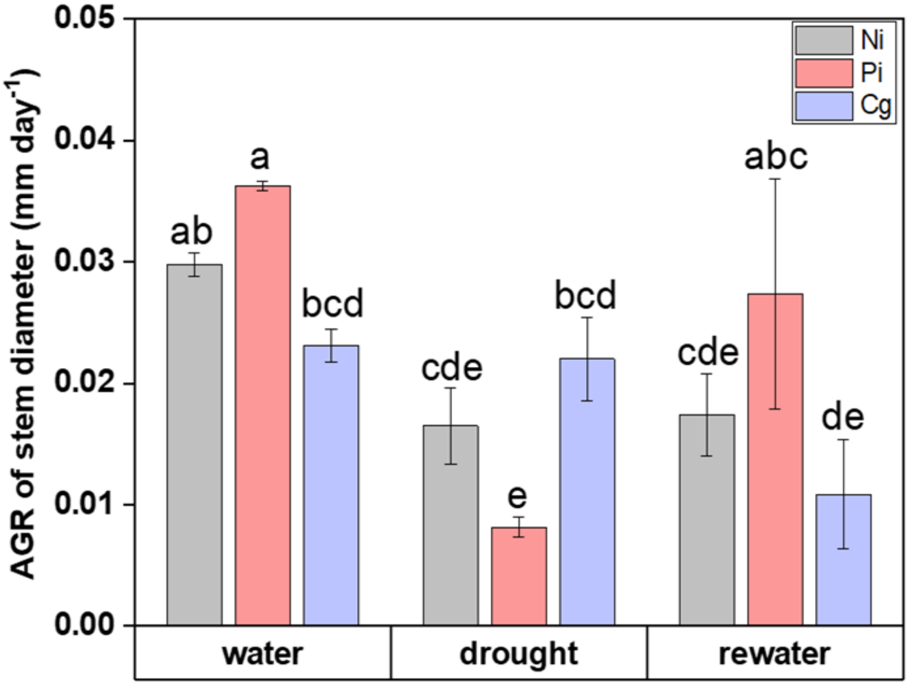


**Supplementary Figure S1**: Diameter growth rate (AGR) of mycorrhizal poplars in response to well-watered, drought and rewatered treatments. Data indicated means ± SE (n = 5 to 6 plant per treatment). Significant differences at $p \leq 0.05$ are indicated by different letters (Two-way ANOVA and post-hoc Fisher's LSD test). Ni: non-inoculated, Pi: *Paxillus involutus*, Cg: *Cenococcum geophilum* Fr., water: well-watered (20 weeks), drought (4 weeks), rewater: re-watered (1 week).

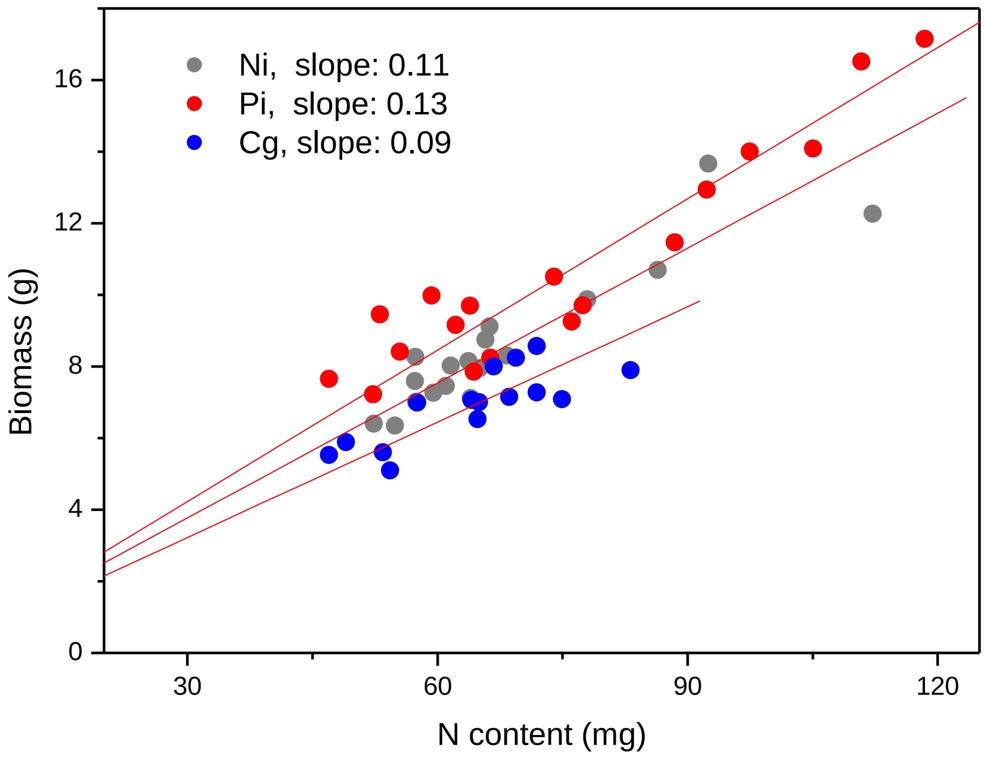


**Supplementary Figure S2:** Relationship between poplar N content and biomass. Linear regression models were applied to poplars colonized with *P. involutus* (Pi), *Cenococcum geophilum* (Cg) and non-inoculated poplars. Each point represents an individual plant. $R^2$adj = 0.9, $p < 0.001$.

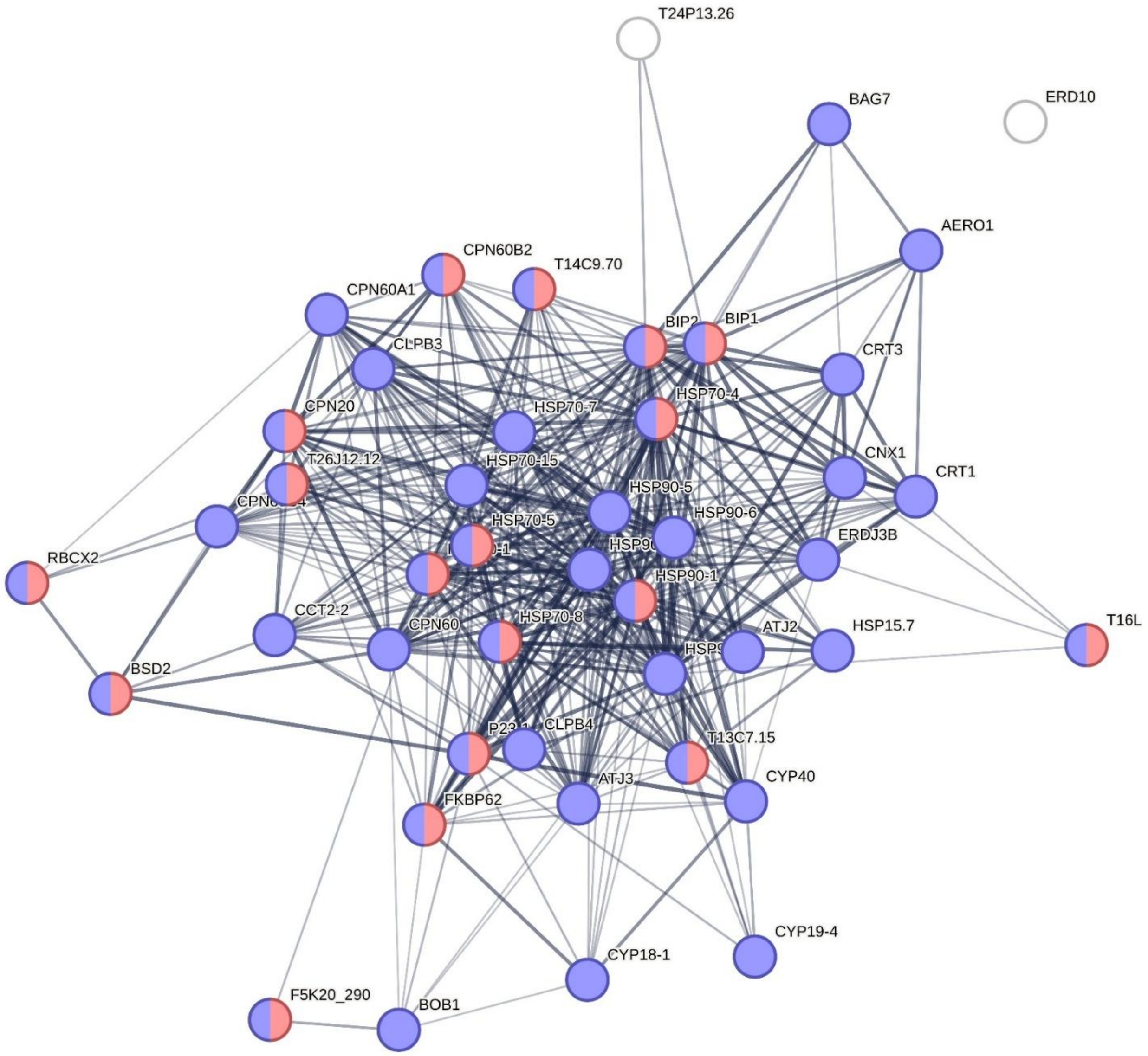


**Supplementary Figure S3**: Network analysis of DEGs in the GO term „Protein Folding“. Blue: protein folding, Red: chaperone mediated protein folding. A total of 45 gene were identified.Number of edges:429, average node degree: 19.1, avg. local clustering coefficient: 0.703, expected number of edges: 25, PPI enrichment p-value: < 1.0e-16
